# Predicting the pathogenicity of missense variants using features derived from AlphaFold2

**DOI:** 10.1101/2022.03.05.483091

**Authors:** Axel Schmidt, Sebastian Röner, Karola Mai, Hannah Klinkhammer, Martin Kircher, Kerstin U. Ludwig

## Abstract

Each individual genome harbors multiple missense variants, which can be systematically identified via genome or exome sequencing. This class of genetic variation can alter the functional properties of the respective protein, and thereby lead to clinically relevant phenotypes, such as cancer or Mendelian diseases. Despite advances in computational prediction scores, the classification of missense variants as clinically significant or benign remains a major challenge. Recently, the structure of the human proteome was derived with unprecedented accuracy using the artificial intelligence system AlphaFold2. However, the question of whether AlphaFold2 structures can improve the accuracy of computational pathogenicity prediction for missense variants remains unclear. To address this, we first engineered a set of features for each amino acid from these structures. We then trained a random forest to distinguish between proxy-benign and proxy-pathogenic missense variants derived from gnomAD. This yielded a novel AlphaFold2-based pathogenicity prediction score, termed AlphScore. Important feature classes used by AlphScore are solvent accessibility, amino acid network related features, features describing the physicochemical environment, and AlphaFold2’s quality parameter (pLDDT). AlphScore alone showed lower performance than existing scores, such as CADD or REVEL. However, when AlphScore was added to those scores, the performance always increased, as measured by the approximation of deep mutational scan data, as well as the prediction of expert-curated missense variants from the ClinVar database. Overall, our data indicate that the integration of AlphaFold2 predicted structures can improve pathogenicity prediction of missense variants.

## INTRODUCTION

The systematic assessment of genetic variation using whole exome or whole genome sequencing (WES/WGS) to identify causal disease variants in individual patients is now an increasingly widespread approach in clinical medicine and medical research. However, this process generates a vast number of novel genetic variants, which require robust annotation of their functional (and potentially pathogenic) effects. While loss-of-function (LoF) variants are relatively easy to interpret due to their immediate deleterious effects, the impact on protein function of missense variants is far less predictable, and their interpretation is therefore problematic. This is exacerbated by the fact that missense variants occur more frequently in individual genomes than LoF variants (Karczewski et al., 2020), and missense variants cover a broad spectrum of functional consequences – ranging from benign to pathogenic - with most being benign (Zuk et al., 2014). Thus in the absence of clear candidate genes or segregation data, identifying potential pathogenic missense variants in the genetic data of any given individual is challenging. To facilitate the assignment of functional effects for missense variants, three different but complementary approaches are generally applied individually or in concert.

In the first approach, analysts access databases containing information on: (i) the frequency of variants in the general population; (ii) the tolerance of genes towards specific types of mutations; or (iii) individual variants that have already been observed and deemed causal in other patients, either by clinical laboratories or on the basis of functional evidence (e.g., ClinVar (Landrum et al., 2018)). In the case of Mendelian diseases, for example, analysts often focus on rare variants in the general population, since these are typically causal (Richards et al., 2015). However, many of the rare missense variants that are observed in WES/WGS data are either novel, or show conflicting entries across databases.

In the second approach, analysts assess the pathogenicity of missense variants using computational prediction scores. In practice, this is a frequently used method, although caution is required due to limited accuracy (Richards et al., 2015). One popular prediction score is Combined Annotation Dependent Depletion (CADD (Rentzsch et al., 2019)). This integrates a wide range of features, such as the missense prediction scores SIFT (Ng and Henikoff, 2003) and PolyPhen-2 (Adzhubei et al., 2010). Recently, two further computational prediction scores have become increasingly popular: 1) REVEL, an ensemble method that combines 13 established prediction tools (including PolyPhen-2 and SIFT) in order to yield a missense variant specific score (Ioannidis et al., 2016); and 2) DEOGEN2 (Raimondi et al., 2017), which includes information concerning the gene, as well as its protein domains and interactions to also yield a missense variant specific score. Benchmarking the accuracy of computational prediction scores is challenging, as independent test datasets are difficult to obtain. Missense variants from ClinVar are potentially biased, since in many cases, computational prediction scores are also used as part of the clinical pathogenicity assessment (Richards et al., 2015). In addition, some existing scores, such as REVEL, use ClinVar variants as training sets, and therefore at least partially recapitulate ClinVar’s ascertainment biases (Ioannidis et al., 2016).

The third approach to missense variant annotation is the performance of deep mutational scans (DMS), which generate experimental data. This approach is becoming increasingly popular in medical research. In DMS, the effect of amino acid substitutions on specific protein characteristics (e.g., stability and folding) or functions (e.g., survival, abundance, binding, metabolic products) are measured via the generation of large-scale datasets, and the results are made available to the research community as independent functional data on the pathogenicity of missense variants (Findlay, 2021). In addition to the annotation of missense variants, DMS also represent a unique resource for the benchmarking of prediction scores for missense variants (Gray et al., 2018; Livesey and Marsh, 2020; Reeb et al., 2020). For instance, research into pathogenicity prediction for missense variants has shown that both REVEL and DEOGEN2 predictions correlate well with DMS data (Livesey and Marsh, 2020), while REVEL also perfomed well on ClinVar data (Tian et al., 2019).

A potentially very useful information level in terms of predicting the pathogenicity of missense variants is the integration of structural data of proteins. To date, information concerning 3D protein structure is sparsely used by missense prediction tools, this may be attributable, in part, to the fact that for ~80% of residues no experimental structures are yet available (Diwan et al., 2021; Somody et al., 2017). However, one of the 11 features used by DEOGEN2 is protein-protein interaction, as detected in experimental protein structures. For proteins with known 3D structures, PolyPhen-2 uses the features accessible surface area of the residue, change in hydrophobic propensity, and the crystallographic B-factor. Recently, the artificial intelligence system AlphaFold2 generated a highly accurate prediction of nearly all 3D protein structures of the human proteom (Jumper et al., 2021; Tunyasuvunakool et al., 2021). AlphaFold2 structures (7,020 of the 16,533 structures used) were already used to analyze constraints regarding missense variation in spatially adjacent amino acids (Li et al., 2021). However, the potential of AlphaFold2-derived protein structures for predicting the pathogenicity of missense variants is, to our knowledge, currently not fully explored.

The aim of the present study was to determine whether the structure predictions of AlphaFold2 can facilitate systematic prediction of the pathogenicity of missense variants. For this purpose, we extracted a set of features from the predicted structures to train tree-based machine learning classifiers. Using variants from DMS and ClinVar, we show that AlphaFold2 structures contain information that is useful in terms of pathogenicity assessment, and that this information can be integrated with existing prediction scores in order to increase their predictive value.

## RESULTS

### Generation of AlphScore

To determine the utility of AlphaFold2 structures for systematical prediction of the pathogenicity of missense variants, we followed the workflow shown in Figure 1. First, structural features were extracted for each amino acid contained in AlphaFold2-predicted structures. To provide a comprehensive description of the environment of each amino acid in the structural models of AlphaFold2, 218 features such as molecular interactions, solvent accessibility, secondary structure, and amino acid network features were extracted (Table S1). Second, the structural features of each amino acid were added to the dbNSFP database (version 4.2), which contains all potential non-synonymous single-nucleotide variants (SNVs) with extensive annotations (Liu et al., 2020). In total, structural features from 17,880 proteins were mapped to dbNSFP (listed in Table S2). Third, three variant sets were created for training, validation, and testing, respectively, using the gnomAD and ClinVar databases (see Methods). GnomAD variants were used as a training set (gnomAD_train, n=315,876), since these variants are assumed to be less prone to biases such as curation, research history, clinical evaluation criteria, and existing computational missense prediction scores (Shah et al., 2018). GnomAD singleton variants and variants with a minor allele frequency (MAF) > 0.1% were used as proxy-pathogenic and proxy-benign variants, respectively. However, independent sets of (probably) benign or (probably) pathogenic ClinVar missense variants were used for validation (ClinVar_val, total n=35,640) and testing (ClinVar_test, total n=21,068).

**Figure 1.**
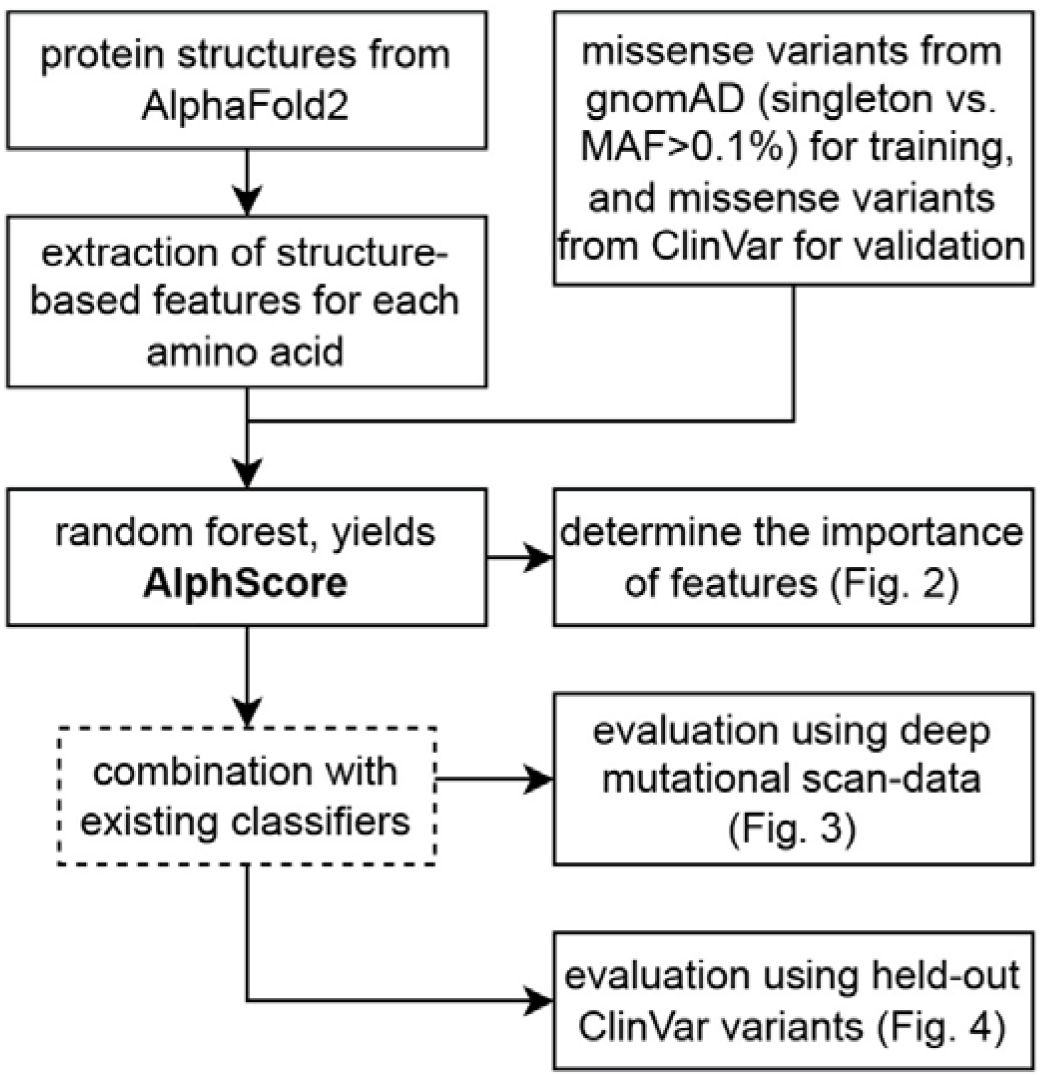
Workflow. MAF: Minor allele frequency.

The gnomAD_train and the ClinVar_val sets of variants were used to train and evaluate three tree-based machine learning algorithms (i.e., gradient boosting, random forest, and extremely randomized trees). Hyperparameters were varied in a grid search. A model using random forests, termed AlphScore, achieved the best overall performance (Area Under the Receiver Operating Characteristics (AUROC) of 0.793 on ClinVar_val, see Table S3). To verify that AlphaFold2-based features play a genuine role in variant classification, the same algorithm was refitted following the removal of AlphaFold2-related features. This model (NullModel) retained only the reference and alternative amino acid as well as simple physicochemical properties of amino acids, and obtained a substantially lower AUROC (0.609). This demonstrates that AlphaFold2-based features are an important component of AlphScore. In the same analysis, CADD achieved an AUROC of 0.871. However, AlphScore relies purely on AlphaFold2-based features, and does not contain features such as amino acid or nucleotide conservation.

### Determination of most important features

To investigate which AlphaFold2-derived features were most important to AlphScore, permutation based feature importances were calculated (Figure 2 and Table S4). Among the top 25 features, the most important AlphaFold2-based feature categories were: solvent accessibility (containing n=8 features); amino acid network related features (n=4); features describing the physicochemical environment (n=4); and AlphaFold2’s parameter for the reliability of its structural predictions, predicted Local Distance Difference Test (pLDDT) (n=2). The top 25 features included seven that were not derived from the structural models of AlphaFold2. Instead, these seven features described the identity or general properties of the reference or alternative amino acid.

**Figure 2:**
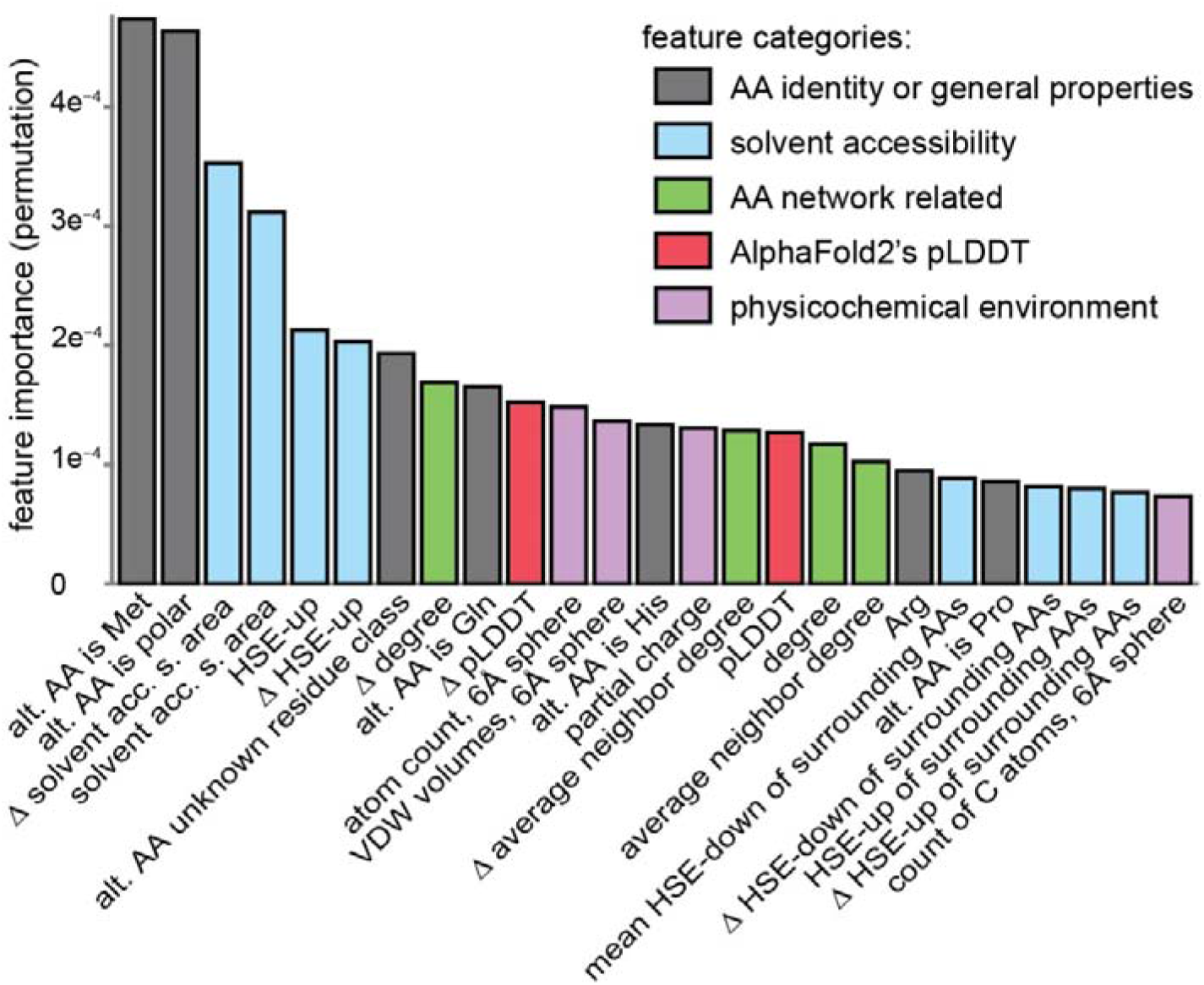
Importance of features in AlphScore. Bar graph displaying permutation-based feature importance for the 25 most important features (total number of features = 218). The feature categories are color-coded. The delta sign represents the difference between the alternative amino acid and the reference amino acid for a given feature (see Methods). Alt. AA: Alternative amino acid. AA: Amino acid. Solvent acc. s. area: Solvent accessible surface area. VDW: Van der Waals. HSE-up: Half-sphere exposure of the upper sphere. HSE-down: Half-sphere exposure of the lower sphere. pLDDT: predicted Local Distance Difference Test, AlphaFold2 per-residue confidence score.

Since the AlphaFold2 pLDDT is an interesting potential new parameter for predicting the pathogenicity of missense variants, pLDDT-based features were removed from the model on a test basis. As a result, the performance on the validation set (ClinVar_val) decreased marginally, from 0.793 to 0.790. Nevertheless, it seems interesting to note that both proxy-pathogenic gnomAD and (likely) pathogenic ClinVar variants tend to have higher pLDDT values (see Figure S3).

### Performance evaluation for AlphScore

To test AlphScore’s performance, AlphScore and the three *in silico* prediction scores were applied to missense variants from DMS or missense variants classified in ClinVar (ClinVar_test). In addition, combined scores of AlphScore and CADD, REVEL, and DEOGEN2, respectively, were created. To determine the best performing parameters for combination, logistic regression was used to fit ClinVar_val variants to combinations of scores (see Methods).

DMS datasets that had been used in prior benchmarking analyses were used (Livesey and Marsh, 2020), and recent data on MSH2 and VKOR1 were added (Chiasson et al., 2020; Jia et al., 2021). In total, the present DMS dataset comprised 13 experiments in 11 proteins (ADRB2, BRCA1, HRAS, MSH2, P53, PTEN, SUMO1, TPK1, TPMT, UBE2I, and VKOR1). Calculations were then performed to determine the Spearman correlations between different computational prediction scores, and combinations thereof, and DMS scores. The average absolute Spearman correlation between AlphScore and the DMS scores was 0.344. In comparison, the established scores achieved an average absolute Spearman correlation with the DMS scores of between 0.338 and 0.422 (Figure 3A). For each prediction score, the addition of AlphScore increased the average Spearman correlation (CADD: 0.338 to 0.399; DEOGEN2: 0.422 to 0.436; REVEL: 0.421 to 0.442). Notably, while the highest overall Spearman correlation with the DMS data (0.450) was achieved by a combination of AlphScore with both DEOGEN2 and REVEL, it was not reached for the combination of the three existing prediction scores alone (Figure 3). To investigate the reliability of those results, we applied 1000-fold bootstrapping and calculation of the rank score, which was introduced by Livesey and Marsh, 2020 (Figure 3B). This rank score is calculated as average across proteins with a scale from 0 to 1. Prediction scores with higher correlations to DMS data receive a value closer to 1. Adding AlphScore improved the rank score for both CADD and REVEL in 1000 and for DEOGEN2 in 950 out of 1000 bootstrapped samples. The combination AlphScore with both DEOGEN2 and REVEL achieved the highest rank score in 1000 out of 1000 bootstrapped samples.

**Figure 3:**
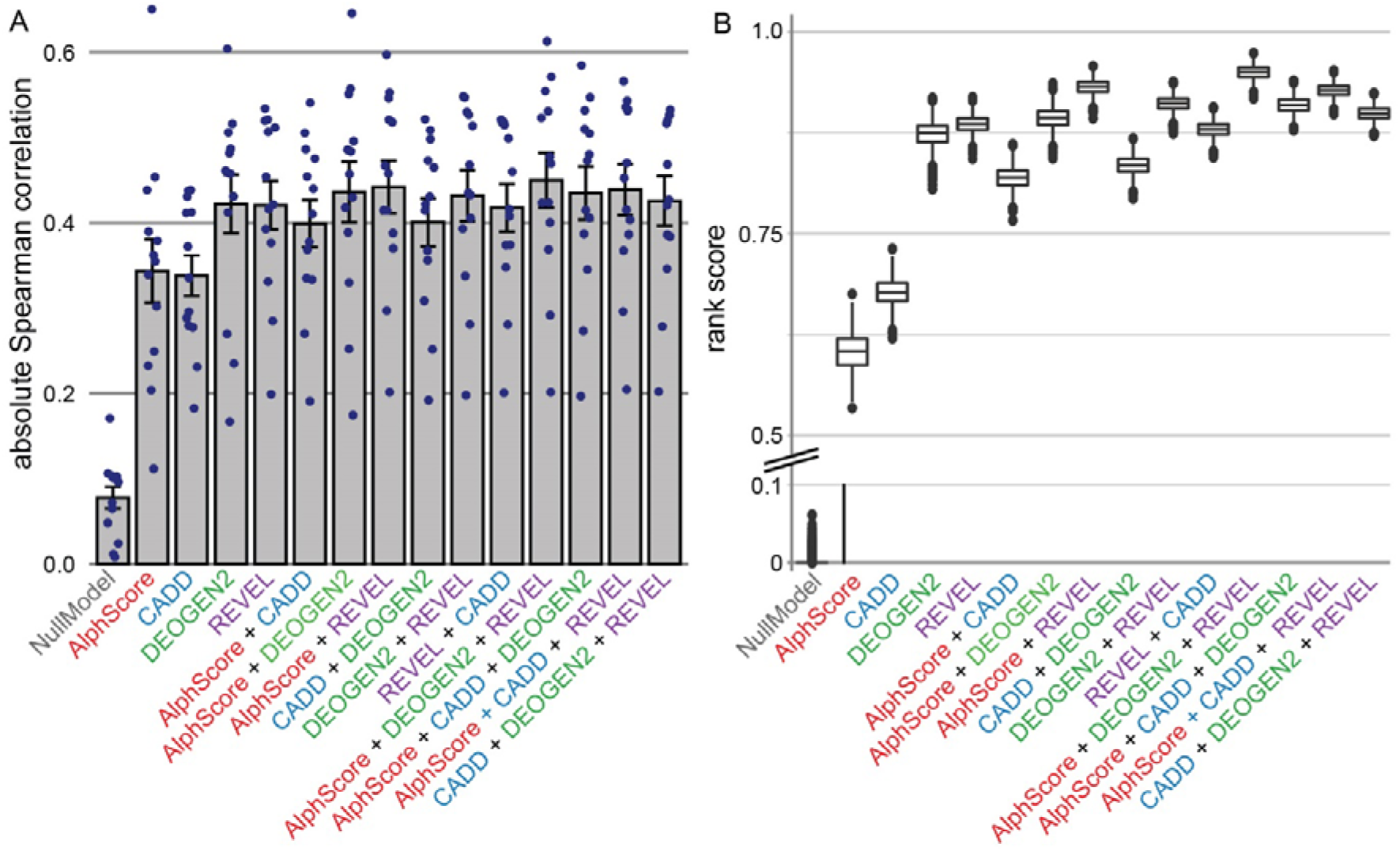
Combining AlphScore with established missense prediction scores improves correlation with deep mutational scans (DMS) data. A) Bar graphs displaying the mean absolute Spearman correlation between DMS scores and computational prediction scores. The dots represent the Spearman correlations of individual DMS experiments. The error bars represent the standard error of the mean. B) Boxplots of rank scores as defined by Livesey and Marsh, 2020, for 1000 bootstrapped samples. Higher rank scores indicate better Spearman correlations with DMS datasets compared to the competing prediction scores. The y-axis has been split to improve visibility. Note that variants in the sets gnomAD_training and ClinVar_val were removed from the analyses.

Investigations were then performed to determine how AlphScore and combinations of AlphScore with existing prediction scores would perform in the test set of ClinVar variants (ClinVar_test, n=21,068), which contained proteins and variants that were independent from those contained in gnomAD_train and ClinVar_val (see Methods). The AUROC of AlphScore alone was 0.799, whereas CADD, DEOGEN2, and REVEL achieved 0.885, 0.886, and 0.929, respectively. Again, in each case, the addition of AlphScore to the established scores increased the respective AUROC (CADD: 0.885 to 0.909; DEOGEN2: 0.886 to 0.907; and REVEL: 0.929 to 0.935; see Figure 4). Use of the Area Under the Curve in precision-recall curves as an alternative measure generated very similar results (Figure S4). Reassuringly, these results could be reproduced in 1000 of 1000 bootstrapped samples of variants.

**Figure 4:**
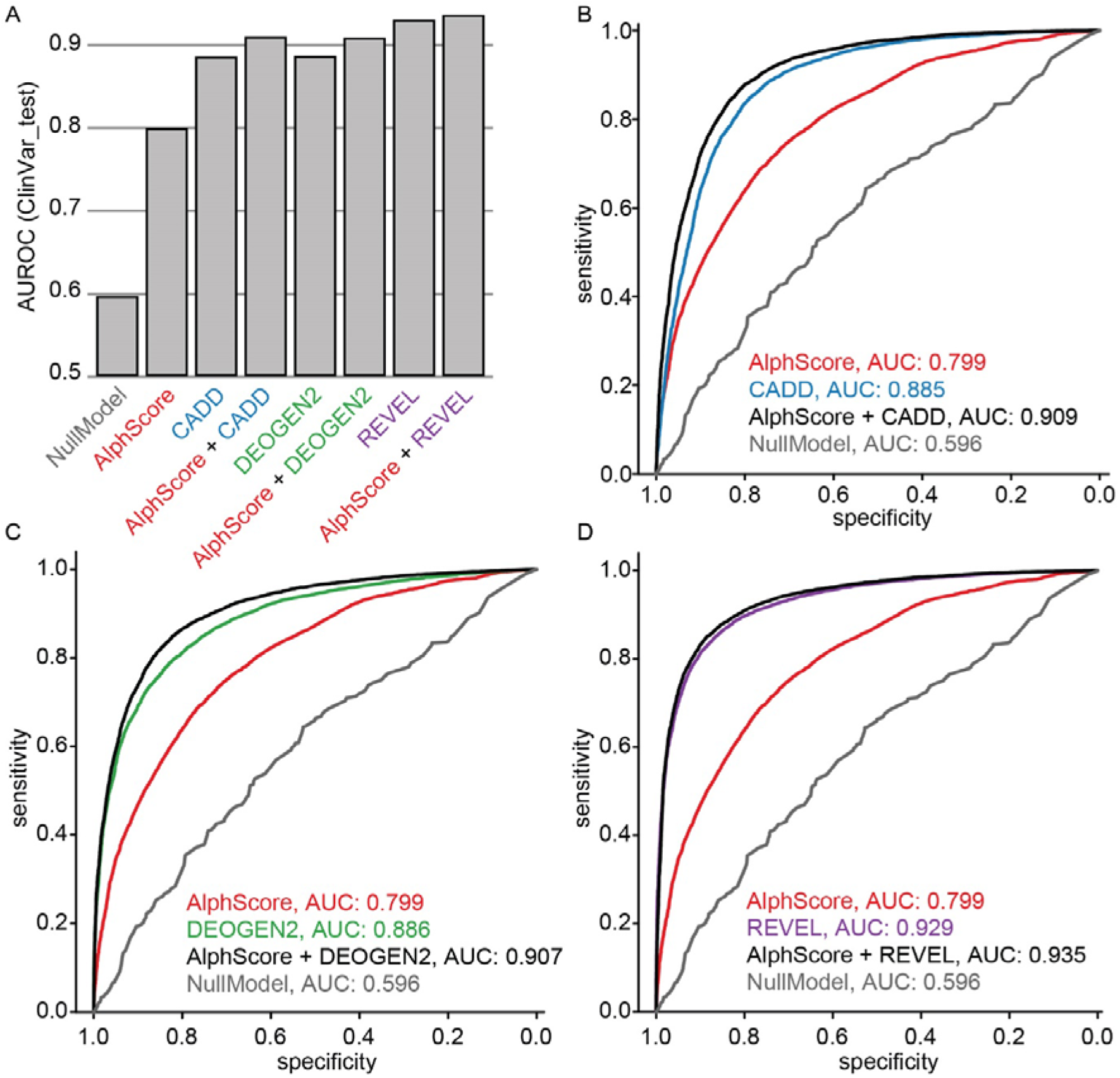
Combining AlphScore and established prediction scores improves the prediction of ClinVar variants. (A) Bar graphs representing the average AUROCs of the prediction scores denoted on the x-axis as obtained from the Receiver Operating Characteristics (ROC) curves shown in B-D. (B-D) ROC curves showing the performance of AlphScore (B-D, red), CADD (B, blue), DEOGEN2 (C, green), REVEL (D, lilac) and linear combination of the AlphaFold-based score with the three existing prediction scores (black). A hold-out set of 9,224 (likely) benign and 11,844 (likely) pathogenic missense variants from ClinVar (ClinVar_test) was used as data source. The gray line (NullModel) represents the baseline model, which was trained on the gnomAD training set, without AlphaFold-based features. AUC: Area Under the Curve.

### Availability of pre-computed scores

Finally, the random forest was retrained with the aim to maximize power. Therefore, no variants in specific proteins were held-out for subsequent testing. The resultant score was termed AlphScore_final. This could be calculated for 80% (n=66,931,527) of dbNSFP’s variants. AlphScore_final and combinations of AlphScore_final with CADD, REVEL or DEOGEN2 are available for download (DOI: 10.5281/zenodo.6288139; Figures S5 and S6 for additional data on the final scores).

## DISCUSSION

The present study generated AlphScore, a novel prediction score for missense variants which relies solely on features derived from the structural predictions of AlphaFold2. Comparisons with experimental high-throughput data and clinically-informed variants from ClinVar showed that the addition of AlphScore improved the performance of established *in silico* missense prediction scores. This suggests that AlphScore captures information that is not encompassed in the existing scores, or that was underweighted in their training. To facilitate future work, the end-to-end pipeline and precomputed scores for 67 million missense variants have been made accessible to the wider research community.

When we analyzed structure-derived feature categories, we identified solvent accessibility as most important feature category for AlphScore. This is in concordance with expectations, since the inaccessible core of a protein is strongly conserved (Overington et al., 1992), and disease associated missense variants are enriched in residues with low solvent accessibility (Savojardo et al., 2020). Furthermore, among the 25 most important features we identified an interesting and easily accessible new parameter, i.e., the quality score of AlphaFold2, pLDDT. Regions with low pLDDT displayed a depletion of both pathogenic variants from ClinVar, and proxy-pathogenic variants from gnomAD. Given that low pLDDT values may be associated with intrinsically disordered protein regions (Ruff and Pappu, 2021; Tunyasuvunakool et al., 2021), this could indicate that these regions tend to be less disease-relevant in aggregate than well-structured regions. However, intrinsically disordered regions are also implicated in important functions, such as mediating interactions with protein domains or harboring posttranslational modifications (Babu, 2016). In general, they also show lower conservation, as well as an altered amino acid composition, in comparison to well-structured regions (Brown et al., 2010), which could partially explain their depletion of pathogenic variants in ClinVar.

Despite the potential of AlphScore to improve existing missense prediction scores, several limitations in the current implementation of AlphScore remain. For instance, AlphScore only considers features that are represented in the structural environment of an amino acid in AlphaFold2 models. Thus, structural features such as post-translational modifications, or features of the amino acid environment resulting from the quaternary structure, are not considered. Furthermore, since AlphScore is based solely on AlphaFold2 structural models, non-coding effects, such as alterations in splicing, are not considered either. Another limitation are proteins that are currently not covered by AlphScore, due to difficulties in mapping between Uniprot and genomic positions (https://www.uniprot.org/help/canonical_nucleotide).

To improve AlphScore, a number of avenues could be explored in future research. First, the choice of extracted features could be refined, or machine learning algorithms that do not require manual feature engineering, such as 3D convolutional neural networks (Torng and Altman, 2017), could be applied. The fact that all proteins of AlphaFold2 were modeled as monomers without ligands and output in a common data format further facilitates the application of such machine learning approaches. Second, the training set could be optimized, e.g., by using variants with stronger signals of purifying selection (such as simulated *de novo* variants as opposed to proxy-pathogenic singleton variants (Kircher et al., 2014)). Finally, additional layers of information, like existing missense prediction scores, could be integrated into the score at an earlier stage, for example, prior to the training of the random forest model. It will be interesting to see the relative importance of structure-based features in such integrated scores. This is particularly true if advanced models of sequence evolution, such as the recently published model EVE, are integrated with structural features, since AlphaFold2 structures themselves are based primarily on sequence evolution data. However, currently EVE only supports a limited set of genes (Frazer et al., 2021). Our opinion is that the implicit or explicit representation of human proteins as 3D structures will remain an important layer of information for missense variant interpretation.

The present study focused on predicting the pathogenicity of missense variants, rather than determining the pathogenic mechanism of a single variant. In principle, to investigate the pathogenic mechanism, allowing the AlphaFold2 model to predict the structure of the protein with the mutant amino acid sequence would be a logical approach. However, research has shown that AlphaFold2 does not accurately predict the structural effect of variants that are known to lead to a structural change (Buel and Walters, 2022). At present, established methods such as molecular modeling should rather be considered to answer these questions. For such methods, and thus for the study of the pathogenic mechanism of missense variants, the AlphaFold2-based structures are a valuable resource.

In summary, we applied a machine learning approach with classical feature extraction to the protein structures generated by AlphaFold2 and demonstrated that these structures contain information that is valuable in terms of predicting the pathogenicity of missense variants. We are eager to see how other groups will use AlphaFold2-based structures to predict the pathogenicity of missense variants and whether approaches, such as modifications of the AlphaFold2 model, might be competitive in predicting the pathogenicity of missense variants.

## METHODS

### Software

The present data were generated using a custom Snakemake pipeline, which is available via github (https://github.com/Ax-Sch/AlphScore). The main frameworks used were Snakemake version 6.12.3, Python version 3.10, and R version 4.1.1.

### Datasets

For training and testing, two missense variant datasets were defined. The first contained missense variants derived from the population database gnomAD (version 3.1 as included in dbNSFP). Variants in this dataset with an allele frequency > 0.1 % were labeled as proxy-benign. In contrast, proxy-pathogenic variants were defined as singleton variants in gnomAD genomes that originated from the non-finish european (NFE) subcohort and which were absent in both the 1000 genomes project and the NHLBI Exome Sequencing Project (ESP6500). In gnomAD exomes, an allele count of 0 or 1 was tolerated for proxy-pathogenic variants. However, an equal allele count in the total cohort and in the NFE subcohort was required. To reduce biases in the training set, the remaining gnomAD variants were subsampled to yield a constant ratio of proxy-pathogenic and proxy-benign variants for each reference amino acid (see Figure S2).

The second dataset was created by filtering ClinVar (version 20210131 as included in dbNSFP) for missense variants labeled as benign/likely benign or pathogenic/likely pathogenic.

Next, 80 percent of proteins were selected at random. The gnomAD variants within these proteins served as the training set (gnomAD_train), whereas variants that were present in ClinVar but not in gnomAD_train served as the validation set (ClinVar_val). ClinVar variants within the remaining 20% of proteins, and ClinVar variants that were new to ClinVar version 20220109, were used as the final test set (ClinVar_test). The independence of the datasets was confirmed by mutual exclusion of variants via chromosome, position, reference allele and alternative allele (see Figure S1).

### Feature extraction

Protein structures predicted by AlphaFold2 for the human reference proteome were downloaded from the EMBL-EBI website (https://alphafold.ebi.ac.uk/download, 29 November 2021). Several tools were applied for feature extraction. First, secondary structures and solvent-accessible surface areas were extracted using DSSP (version 3.0.0, (Kabsch and Sander, 1983)). Second, the FEATURE framework (version 3.1.0, (Halperin et al., 2008)) was applied to calculate physicochemical features within spheres with diameters of 0, 3, and 6 angstroms. The values of the C-alpha atom of the FEATURE framework were selected for each residue. Inter-residue interactions and contacts were also extracted using a modified version of the Protinter software (https://github.com/Ax-Sch/protinter). Half-sphere exposures and the pLDDT were extracted using the Biopython or the Biopandas python package, respectively. Finally, a weighted amino acid network was constructed for each protein using the python package biographs and an atom-atom distance cutoff of 4 angstroms (https://github.com/rodogi/biographs). This network was used to calculate weighted means of the properties of neighbor amino acids, as well as network based metrices, using the python package networkx (see Table S1). In cases where several structural models were available for one protein, the average value for each feature across the structural models was used. The features obtained for each amino acid were then added to the dbNSFP4.2a database using the UniProt ID and the Variant Effect Predictor annotations as keys (Liu et al., 2011; Liu et al., 2020).

### Data preprocessing

In addition to the features described above, 20 binary variables encoding the reference amino acid and 20 binary variables representing the alternative amino acid were created (see Table S1). To optimize representation of the effects of amino acid substitutions, the average values of selected features for each of the 20 possible amino acids were determined. For this purpose, the gnomAD_train dataset was used and filtered for residues with high confidence prediction (pLDDT>90). The average value obtained was attributed to the alternative amino acid for each missense variant. The difference between this average value assigned to the alternative amino acid and the value extracted for the reference amino acid was then calculated. This difference was appended to the dataset.

### Machine learning and grid search

Due to the robustness and interpretability of tree-based predictors, gradient boosting (R package xgboost), random forests (R package ranger), and extremely randomized trees (R package ranger) were selected as candidate algorithms. Several algorithm / parameter combinations were evaluated during the grid search, as shown in Table S3. The performance of each model was evaluated in ClinVar_val by calculating the AUROC. The best model (AlphScore) used random forests, as implemented in the package ranger with 2000 trees (num.trees), a maximal depth (max.depth) of 5, and a minimal node size (min.node.size) of 10. Otherwise, default parameters were used. Prior to model fitting, identical features were pruned (correlation higher than 0.999999). The features listed in Table S1 were used for prediction.

### Analysis of model characteristics

To calculate permutation-based feature importance for AlphScore, the option importance=“permutation” was set in the package ranger during model fitting. To assess the relevance of certain groups of features, additional models were fitted. These models used the parameters of the top performing model, with the exception that either all AlphaFold-derived features (NullModel) or all features containing AlphaFold’s pLDDT parameter were removed from the provided predictor variables.

### Combination with other missense prediction scores

AlphScore was combined with established missense prediction scores using logistic regression as implemented in the R-function glm with the option family=binomial(link=‘logit’). The validation dataset ClinVar_val was used as the source of training data.

### DMS data and evaluation

DMS data for the following proteins were taken from the supplementary table provided by Livesey and Marsh, 2020: UBE2I, SUMO1, TPK1 (Weile et al., 2017); BRCA1 (Findlay et al., 2018; Starita et al., 2015); P53 (Giacomelli et al., 2018); HRAS (Bandaru et al., 2017); PTEN (Matreyek et al., 2018; Mighell et al., 2018); and ADRB2 (Jones et al., 2020). DMS data for MSH2 and VKOR1 were added (Chiasson et al., 2020; Jia et al., 2021). In cases where a study had conducted different experiments under different conditions, the condition with the highest overall absolute correlation to the computational predictors was used. When DMS data for <20 variants were available for any computational score, the respective DMS dataset was excluded. This filter led to the exclusion of the protein CALM1 from Weile et al., 2017. To ensure comparability, variants of DMS datasets for which values from one of the prediction scores were missing were excluded. Spearman correlations between the DMS scores and the computational scores were then calculated using the cor.test command within R. In the case of negative correlations, the absolute value was taken. To assess the performance of the scores, we primarily calculated their respective mean absolute Spearman correlation with the 13 DMS datasets. As an alternative measure, we also calculated rank score statistics as previously described in Materials and Methods of Livesey and Marsh, 2020.

### Bootstrapping

To estimate the uncertainty in our results, we used bootstrapping. We generated 1000 bootstrapped samples from the variants for which DMS data were available and from the ClinVar_test dataset, respectively, using the R package boot with default parameters. In each of these samples, we then calculated rank score statistics (DMS data) or AUROCs (ClinVar_val).

### Generation of the final model (AlphScore_final)

The parameters of the best performing prediction model were used to fit a model on the full set of gnomAD variants (not restricted to certain genes). This model was then combined with CADD, REVEL, and DEOGEN2 using logistic regression and all (likely) benign / (likely) pathogenic ClinVar variants that were not contained within the training set (as described above). The model was evaluated with ClinVar variants that were new in ClinVar version 20220109 to ensure that this model had similar properties to AlphScore (see Figure S6 and S7).

### Generation of figures

Figure 1 was created with the software draw.io (https://github.com/jgraph/drawio). Figures were assembled using Adobe Illustrator (version CS6).

## DATA ACCESS

AlphScore_final and combined scores computed in the present study have been submitted to Zenodo and are available under the DOI 10.5281/zenodo.6288139. The computational pipeline used to generate the data presented is available via github (github.com/Ax-Sch/AlphScore).

## ACKNOWLEDGMENTS

AS is supported by the BONFOR program of the Medical Faculty, University of Bonn (O-149.0134). KUL is supported by the Emmy-Noether program of the German Research Foundation (DFG; LU 1944/3-1).

